# Chromosomal rearrangements are commonly post-transcriptionally attenuated in cancer

**DOI:** 10.1101/093369

**Authors:** Emanuel Gonçalves, Athanassios Fragoulis, Luz Garcia-Alonso, Thorsten Cramer, Julio Saez-Rodriguez, Pedro Beltrao

## Abstract

Chromosomal rearrangements, despite being detrimental, are ubiquitous in cancer and often act as driver events. The effect of copy number variations (CNVs) on the cellular proteome of tumours is poorly understood. Therefore, we have analysed recently generated proteogenomic data-sets on 282 tumour samples to investigate the impact of CNVs in the proteome of these cells. We found that CNVs are post-transcriptionally attenuated in 23-33% of proteins with an enrichment for protein complexes. Complex subunits are highly co-regulated and some act as rate-limiting steps of complex assembly, indirectly controlling the abundance of other complex members. We identified 48 such regulatory interactions and experimentally validated AP3B1 and GTF2E2 as controlling subunits. Lastly, we found that a gene-signature of protein attenuation is associated with increased resistance to chaperone and proteasome inhibitors. This study highlights the importance of post-transcriptional mechanisms in cancer which allow cells to cope with their altered genomes.

## Introduction

Cancer development is driven by the acquisition of somatic genetic variation that includes point mutations, copy number variations (CNVs) and large chromosome rearrangements or duplications (i.e aneuploidy) (Beroukhim et al., 2010). These events can result in a fitness advantage and cancer progression but they are most often detrimental to cellular fitness. While somatic gene amplification of key oncogenes such as MYCN, AKT2, ERBB2 and others (Santarius et al., 2010) can drive cancer development, germline CNVs are rare and are under negative selection (Itsara et al., 2009). Gene amplifications and other CNVs are thought to be detrimental due to changes in gene-expression that cause an imbalance to the cell. In females, one of the two X chromosomes is inactivated by a specialized RNA based silencing mechanism (Avner and Heard, 2001; Lyon, 1961) but such mechanism does not exist for gene dosage imbalances in the autosomal chromosomes. Protein and mRNA abundance measurements in models of aneuploidy in yeast (Dephoure et al., 2014) and human cells (Stingele et al., 2012) have shown that most autosomal gene duplications are propagated to the protein level, with the notable exception of protein complex subunits (Dephoure et al., 2014; Stingele et al., 2012). The lack of attenuation of changes in expression resulting from duplicated chromosomes causes global stress responses that include cell-cycle and metabolic defects and proteotoxic stress among others (Tang and Amon, 2013). While somatic CNVs are known to be drivers of cancer development and that aneuploidy is a common feature of tumor cells, the impact of gene dosage changes on the proteome of cancer cells have yet to be studied.

Comprehensive characterisation of hundreds of tumour samples at the genomic and transcript level have been instrumental for the study of tumour heterogeneity across patients and to assess the implications of genomic alterations to cancer development (Cancer Genome Atlas Research Network et al., 2013). Functional annotation of genomic events have been limited to the transcript level until very recently, when the first broad proteomics studies with deep coverage of the proteome were made available (Mertins et al., 2016; Zhang et al., 2014, 2016).

In this study, we investigated the implications of CNVs on the proteome of tumours by taking advantage of the comprehensive data-sets made available by the TCGA and CPTAC consortiums comprising copy-number, transcript and protein measurements for hundreds of tumours (Cancer Genome Atlas Network, 2012a, 2012b; Cancer Genome Atlas Research Network, 2011; Mertins et al., 2016; Zhang et al., 2014, 2016). This data revealed that CNVs are often propagated to the protein level although we observed that post-transcriptional mechanisms attenuate this impact in 23-33% of the measured proteins. Protein complexes were notably attenuated, likely due to the degradation of free subunits, resulting in strong protein abundance coregulation across samples. Not all complex subunits are attenuated, with some acting as potential rate-limiting factors for complex assembly, and we identified 48 novel and known regulatory interactions, whereby the abundance of one of the subunits can modulate the abundance of other complex members. We experimentally validated AP3B1 and GTF2E2 as potential rate-limiting subunits capable of controlling the degradation rate of interacting partners. In addition, ranking the samples by their potential to attenuate gene dosage effects identified putative mechanisms involved in autosomal gene dosage compensation. Finally, a gene expression signature of attenuation potential was found to be associated with drugs targeting chaperones, the proteasome and the E3 ligase MDM2. Using 282 tumour samples we revealed the widespread importance of post-transcriptional mechanisms to ameliorate the impact of CNVs in cancer cells.

## Results

### Tumour pan-cancer proteomics reveals attenuation of copy-number alterations in protein complex subunits

In order to study the implication of gene dosage changes on the proteome of cancer cells we compiled and standardized existing data-sets made available by the TCGA and CPTAC consortiums, comprising 3 different cancer types: breast (BRCA) (Cancer Genome Atlas Network, 2012b; Mertins et al., 2016), high-grade serous ovarian (HGSC) (Cancer Genome Atlas Research Network, 2011; Zhang et al., 2016) and colon and rectal (COREAD) (Cancer Genome Atlas Network, 2012a; Zhang et al., 2016) (Figure 1A). These data-sets provide molecular characterisation of gene CNVs, gene expression and protein abundance of solid tumour samples of 282 patients for which clinical information is also available (Figure 1A, Supp. Table 1).

**Figure 1.**
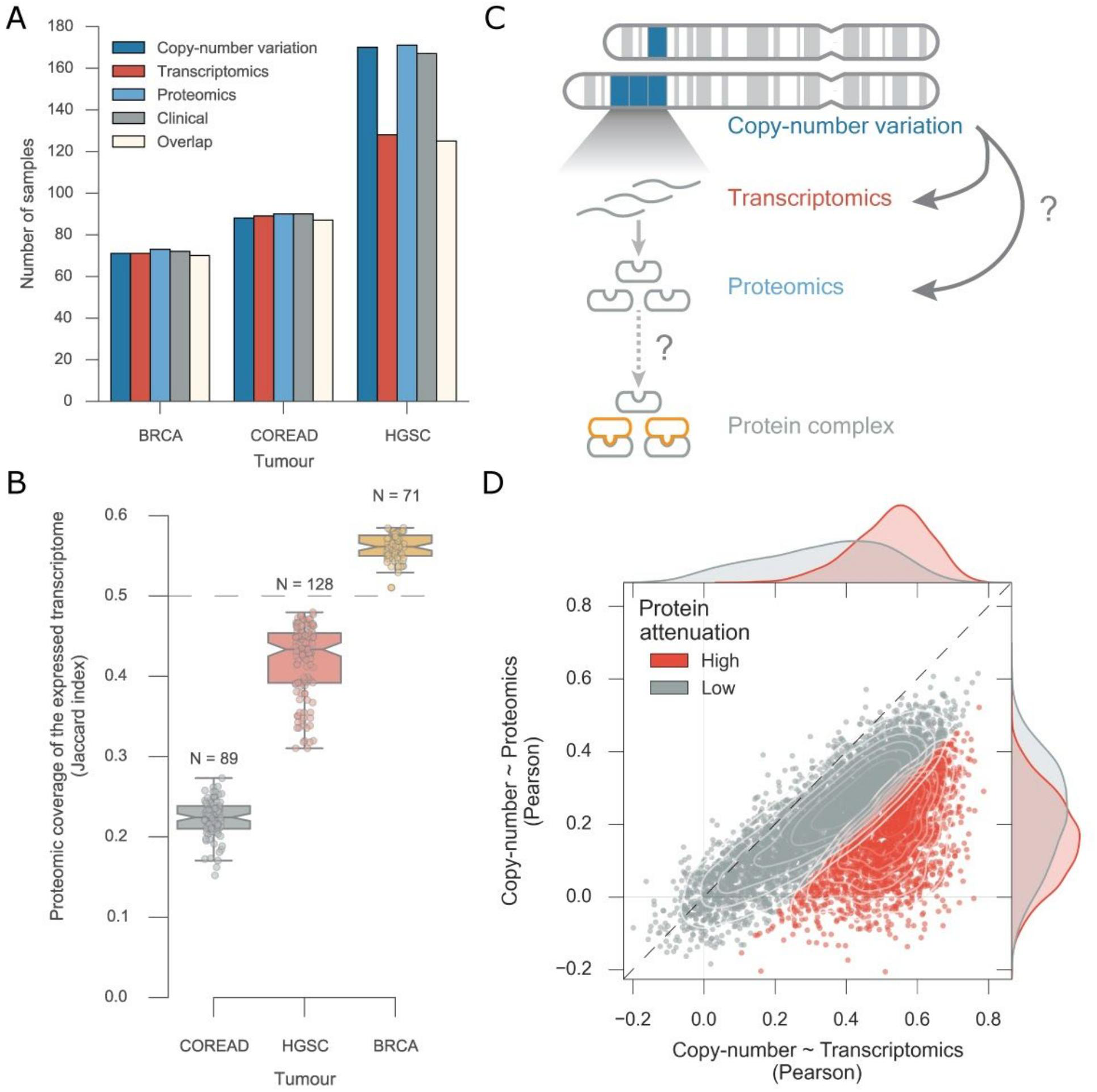
Pan-cancer effects of copy-number variation on transcript and protein abundances. A) Overview of the number of samples used in this study overlapping with the proteomics measurements for each tumour type. B) Proteomics coverage of the expressed transcripts in each sample and for each tumour type. C) Diagram depicting the implication of copy-number alterations along the central dogma of biology. D) Pearson correlation between copy-number variation and transcriptomics in the x axis and copy-number variation and proteomics in the y axis. A gaussian mixture model with two mixture components was used to identify proteins with high attenuation levels (colored in red).

Current methods can reliably measure the complete expressed transcriptome but measuring the total proteome is still a challenge with current techniques only providing partial snapshots (Nagaraj et al., 2011). Thus, we quantified the fraction of expressed transcripts measured in the proteomics experiments in each tumour sample (Figure 1B) (see Methods). COREAD samples displayed the lowest average coverage of the expressed transcriptome (22.3%) compared to the coverage measured for the HGSC (42.0%) and BRCA (56.1%) samples. The proteomics experiments were not conducted using the same methodologies, and therefore it is crucial to take in consideration the potential confounding effects. In particular, the COREAD (Zhang et al., 2014) quantifications were done with a label free approach while the HGSC and BRCA were quantified using isobaric labelling (Mertins et al., 2016; Zhang et al., 2016). To ensure comparable measurements among data-sets we removed confounding and systematic effects from the proteomics and transcriptomics, by regressing-out batch effects associated with experimental technologies used, patient gender and age and tumour type (see Methods). The associations between these possible confounding factors and the principal components were completely removed after correction (Figures S1 and S2).

Having assembled this compendium of data-sets we then set out to understand the implication of CNVs events in the expression of the proteome (Figure 1C). For each gene we calculated the agreement between the CNVs and transcriptomics and CNVs and proteomics using the pearson correlation coefficient (Figure 1D). Transcript abundance is on average well correlated with gene CNVs changes (median pearson’s r=0.43) and this contrasts with the significant decrease (Welch’s t-test p-value < 1e-4) of agreement of CNVs with protein abundance (median pearson’s r=0.20) (Figure 1D, Supp Table 2). As transcription is intermediate between the copy-number alterations and protein abundance, it sets the maximum possible agreement between both. We can then define as attenuated genes those that have a lower agreement between CNVs and protein abundance than expected by their CNV to gene-expression correlation (see Methods). In these samples we found 1,496 - 2,119 genes that are significantly attenuated by this definition, corresponding to 23-33% of all genes with available measurements (6,418). This result shows that a significant fraction of the proteome undergoes gene dosage balancing. Additionally, this group of attenuated genes highlight the complexity of the regulation of protein abundance, hinting at regulatory constraints that control protein translation or degradation rates.

To understand the biological processes that are affected by this attenuation we performed an unbiased enrichment analysis using GO terms (Ashburner et al., 2000; Subramanian et al., 2005; The Gene Ontology Consortium, 2015) (Figure 2A) (see Methods). The enrichment analysis revealed that proteins involved in complexes and modules of functionally interacting proteins displayed a significant agreement at the transcript with the copy-number measurements but this agreement is generally lost at the protein level (Figure 2B). This attenuation of protein complex subunits recapitulates previous findings in models of aneuploidy in yeast (Dephoure et al., 2014) and human cell lines (Stingele et al., 2012) showing that the underlying mechanism generalizes from the aneuploidy models to the hundreds of patient tumour samples studied here. In order to validate the generality of the set of attenuated genes we confirmed that these are also recapitulated in an independent proteomic panel of triple negative breast cancer and ovarian cancer cell lines (Coscia et al., 2016; Lawrence et al., 2015) (Figure 2C). This attenuation mechanism has been shown, in yeast aneuploid strains, to be mostly due to control of protein abundance by degradation (Dephoure et al., 2014). For protein complexes in particular, this result fits with a model where subunits are degraded when free from the complex (Abovich et al., 1985). To test if degradation plays a role in the attenuation observed in human cells, we used publically available data on changes in protein ubiquitination after proteasome inhibition as markers of degradation (Kim et al., 2011). We observed that genes defined as attenuated in our study show a faster increase in ubiquitination after proteasome inhibition than other genes (Figure 2D), suggesting that degradation plays a key a role in the attenuation of these genes. Recently some studies have shown, in degradation time-series experiments, that many protein complex subunits have degradation profiles that are best fit by a two-state model, suggesting that the degradation rate of these proteins changes, presumably when free or when assembled into the complex (McShane et al., 2016). These results suggest that the abundance of protein subunits of large stable protein complexes are under active control to maintain their co-regulation, possibly to guarantee the stability and formation of the associations or prevent the accumulation of free subunits that might be prone to aggregate.

**Figure 2.**
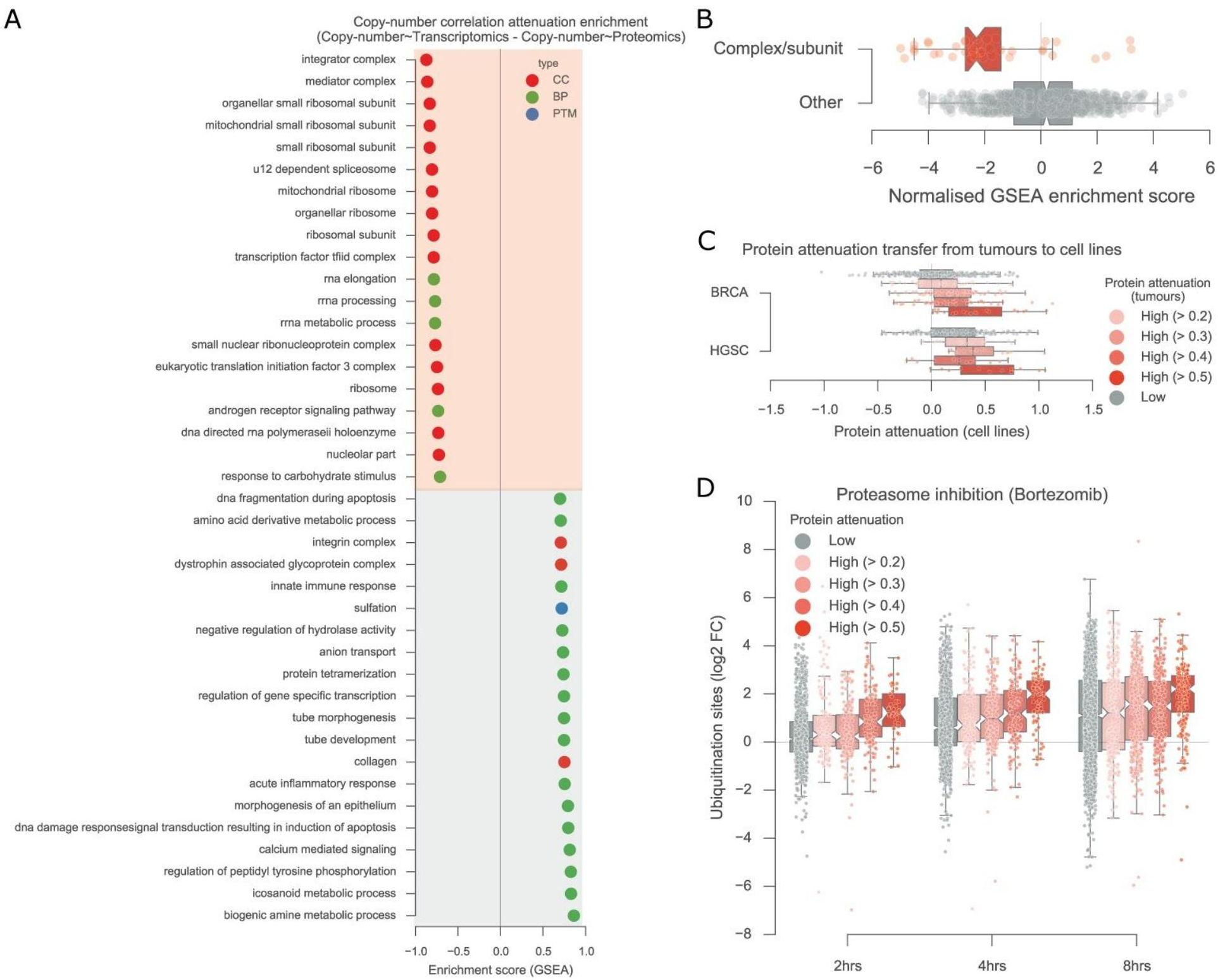
Enrichment analysis of the proteins undergoing copy-number attenuation. A) Enrichment analysis of the correlation differences between copy-number variation and transcriptomics and copy-number variation and proteomics. Protein subsets used represent biological processes (BP; green), cellular components (CC; red) and post-translational modifications (PTM; blue). B) The distribution of the enrichment scores for terms referring to protein complexes or subunits are represented in red and all the rest in gray. C) Proteins classified according to their attenuation profile in tumours are mapped against their attenuation in breast and ovarian cancer cell lines. D) Ubiquitination site fold-changes over time after proteasome inhibition with bortezomib discretized according the protein attenuation level in tumours.

### Proteomic correlation analysis uncovers strong co-regulation of protein complexes

Considering that protein complexes seem to be coregulated to maintain their abundance, likely via the degradation of free subunits, we systematically explored this by performing protein-protein correlation analysis using the proteomics measurements (Figure 3A). We performed all possible pair-wise correlation of protein abundance for all the 6,434 proteins measured in at least 50% of the samples across the 3 different tumour types (see Methods). Consistently, proteins within the same complexes display coordinated changes of abundance across samples (Figure 3A). Then, we assessed if this co-regulation effect is ubiquitous in a curated set of human protein complexes from the CORUM database (Ruepp et al., 2010). Pairs of proteins present together in a complex display a degree of co-regulation (mean pearson’s r=0.25) that is significantly higher than what is observed for random pairs (mean pearson’s r=0). We also assessed if this co-regulation was visible at the transcript level, and while there is a significant increase over random associations (mean pearson’s r=0.15) this correlation is significantly lower than the one seen at the protein level (Figure 3B). Protein pairs that have functional interactions but are not complex subunits show a lower degree of abundance correlation (mean pearson’s r=0.15) that is also closer to the observed at the transcript level (mean pearson’s r=0.11) (Figure 3B).

**Figure 3.**
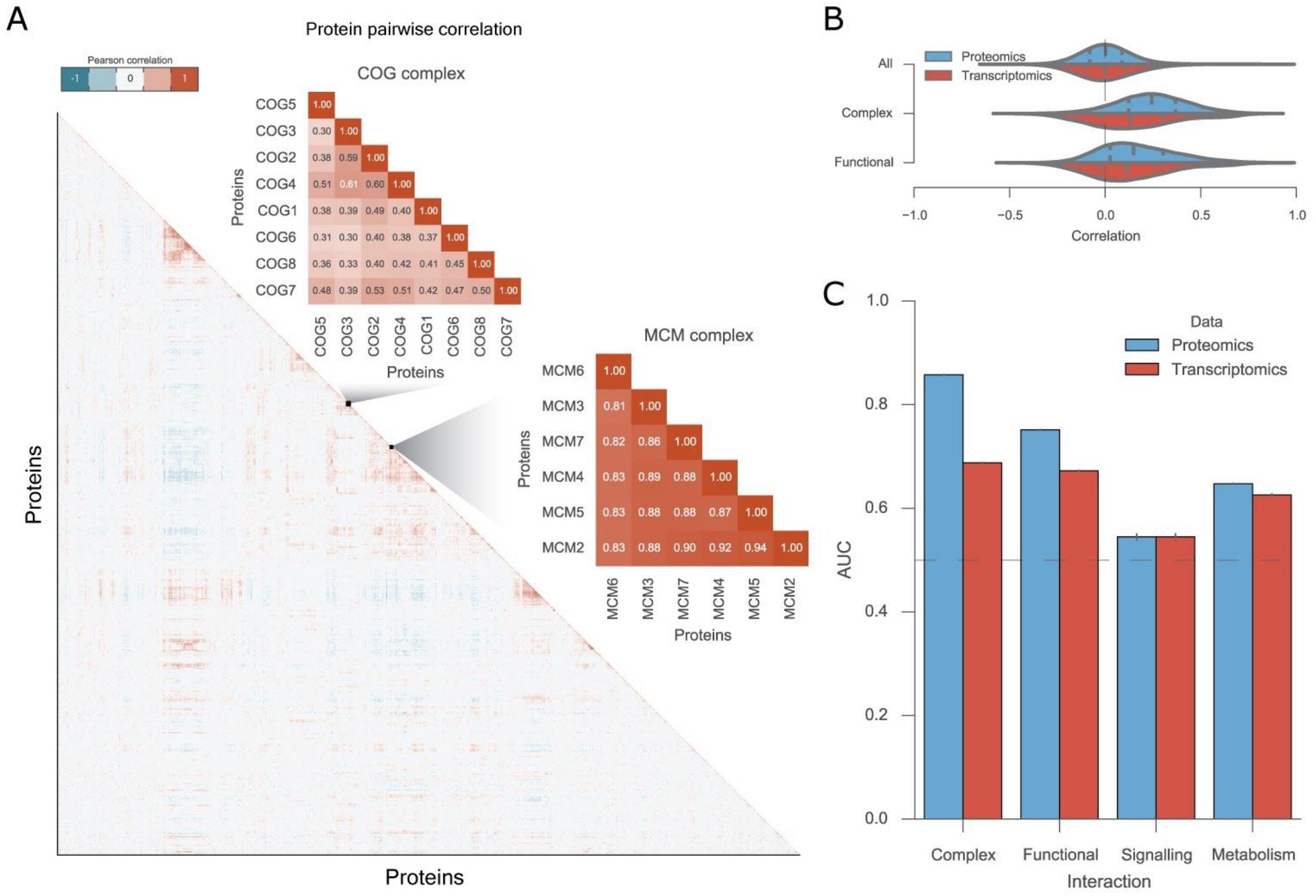
Copy-number variation attenuation for protein complex subunits results in strong co-regulation of their abundances across samples. A) Protein-protein correlation matrix using pearson correlation coefficient and two representative cases of top correlated protein complexes. B) Distribution of all protein-protein correlation at the protein level (Proteomics), and transcript level (Transcriptomics). Protein interactions within complexes are represented by the Complex label and protein functional interactions, that are not necessarily direct, are represented by the Functional label. C) Enrichment analysis by the means of the area under the receiving operating characteristic curves (AROC) using pairwise correlation coefficients, for both proteomics and transcriptomics measurements. Error bars display the variability obtained with five randomised true negative sets.

In light of this agreement between functionally related proteins we examined the capacity of protein-protein correlation profiles to predict known protein-protein interactions (Figure 3C) (see Methods). We found that direct and indirect functional interactions could be well identified with proteomics (AROC 0.86 and 0.75, respectively), and significantly worst predicted with transcriptomics (AROC 0.69 and 0.67, respectively) (Figure 3C). This finding goes in line with a recent work that showed that proteins within similar biological processes or pathways display better agreement at the protein than at the transcript level (Wang et al., 2016). We noticed that protein interactions derived from signalling networks displayed in general poor agreement at the protein and transcript abundance levels (AROC 0.55 and 0.54) (Figure 3C), suggesting that the abundance of signalling proteins in the same pathway does not necessarily need to be coordinated. Furthermore, metabolic enzymes involved in the same metabolic pathways displayed some degree of agreement at the protein and transcript level (AROC 0.65 and 0.62) (Figure 3C).

### Proteogenomics analysis identifies subunits that control the protein abundance levels of other members of the complex

Previous studies conducted in aneuploidy models have indicated that, while protein complex members tend to be attenuated, not all subunits are. It has been hypothesized that these non-attenuated subunits could act as scaffolding proteins or rate-limiting for the assembly of the complex (Dephoure et al., 2014). However, these studies were conducted on a small number of yeast strains or cell lines (Dephoure et al., 2014; Stingele et al., 2012). Given the large number of tumour samples analyzed here we reasoned that we could more readily identify such subunits that can act as drivers of complex assembly. To study this we assessed if CNVs of a given protein within the complex could explain the changes in abundance of other subunits once we discount their transcriptional changes (see Methods). In other words, if the presence or absence of certain proteins of the complex could be associated with the protein degradation rate of other members. This was performed systematically for all identifiable protein-pairs within protein complexes using linear regression models where the CNVs of a protein (Px) was used to estimate the protein abundance variation of the paired protein (Py) (Figure 4A) (see Methods). To consider the differences in degradation or translation rates of the protein, the transcript measurements were regressed-out from the protein abundance (Figure 4A) (see Methods). This allowed us to consider the variability arising only from post-transcriptional regulatory events and, importantly, to discard possible confounding regulatory events occurring at the genomic and transcript level, such as close genomic localization. Out of the 58,627 possible directed protein interactions 64 were found to be significantly associated (FDR < 5%) (Figure 4A, Supp Table 3) (see Methods). To ensure that the association was not only visible at the genomic but also at the transcript level, the same associations were performed using transcriptomics measurements. As expected since that transcript abundance is a closer measurement to the protein abundance, we found a substantial increase of significant associations, 2,846 (FDR < 5%) (Figure S3, Supp Table 3). Also, 75% (48) of the associations found at the genomic level were found to be significant at the transcript level (Figure 4A, Supp Table 3).

**Figure 4.**
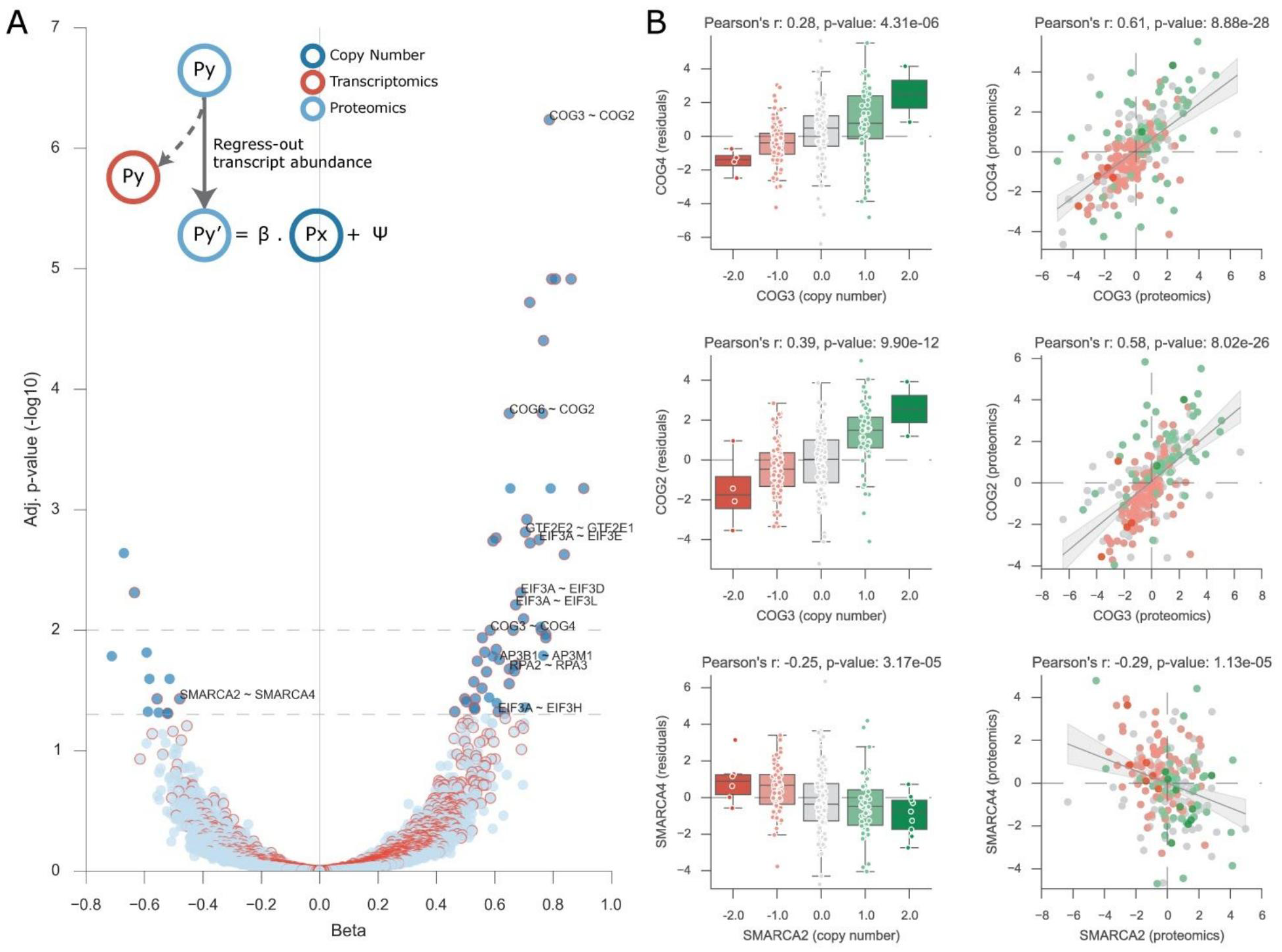
Protein complex regulators. A) Volcano plot displaying the effect size and adjusted p-value of all the tested regulatory interactions. Associations were performed using the copy-number variation of the putative regulatory protein, Px, and the protein residuals of the regulated protein, Py. Significant associations found with the transcript measurements of Px are denoted with a red border. B) Representative significant associations. Boxplots show the agreement between the copy-number variation of Px and the residuals of the regulated Py. Scatter plot show the agreement between the protein pairs in the proteomics measurements.

This analysis identified putative regulators of the assembly of protein complexes that are potentially rate-limiting for the assembly of the full complex. We found, for example, an association between the copy-number of COG3 and the protein variability of COG2 (pearson’s r=0.39, p-value 9.90e-12) (Figure 4B). COG3 is also significantly associated with COG4 (Figure 4B) increasing the possibility that COG3 is a regulator of the assembly of the COG complex. These findings are corroborated by an existing study where COG3 knockdown leads to a decreased abundance of COG2 and COG4 (Zolov and Lupashin, 2005). Besides hinting at known regulatory interactions our analysis also points to possibly novel interactions within the COG complex with COG6 being significantly associated with COG2 (Figure 4A, C). Additional positive regulatory interactions were found for subunits of the eukaryotic Initiation Factor 3 (eIF3), Transcription Factor IIH (TFIIH), Adaptor Related Protein Complex 3 (AP3), among others (Supp Table 3), providing with information on the putative assembly pathways of these complexes.

The number of significant negative associations was remarkably lower than the number of positive associations (Figure 4A, Figure S3 C). SMARCA2 copy-number alterations were significantly negatively associated with the degradation of SMARCA4 (Figure 4A) and this was also visible at the protein level (Figure 4B). Negative associations are likely to represent mutually exclusive events of the protein complex regulation, thus when one protein is present the other will not be necessary for the complex formation and may undergo degradation. Indeed, current evidence in the literature suggest that SMARCA2 and SMARCA4 are paralogs (Ori et al., 2016) and mutually exclusive (Karnezis et al., 2016) within the SWI/SNF complex (Ori et al., 2016). The lower number of negative associations suggests that these types of events are less frequent.

We experimentally validated two of the top significant positive associations (Figure 5A). These were found within protein complex subunits of the adaptor protein complex 3 (AP3) and the Transcription initiation factor IIE (TFIIE), AP3B1 - AP3M1 and GTF2E2 - GTF2E1 respectively. To assess their implication we performed shRNA knockdown of the putative regulatory proteins, AP3B1 and GTF2E2, in shRNA transfected HCT116 human colon cancer cell lines followed by western blot. This showed that knocking down the regulatory proteins not only affected their abundance, AP3B1 and GTF2E2, but also the abundance of the interacting proteins within the protein complex subunit, AP3M1 and GTF2E1 (Figure 5B). This supports the validity of the regulatory interactions found and that these can be transferred between tumours and cell lines.

**Figure 5.**
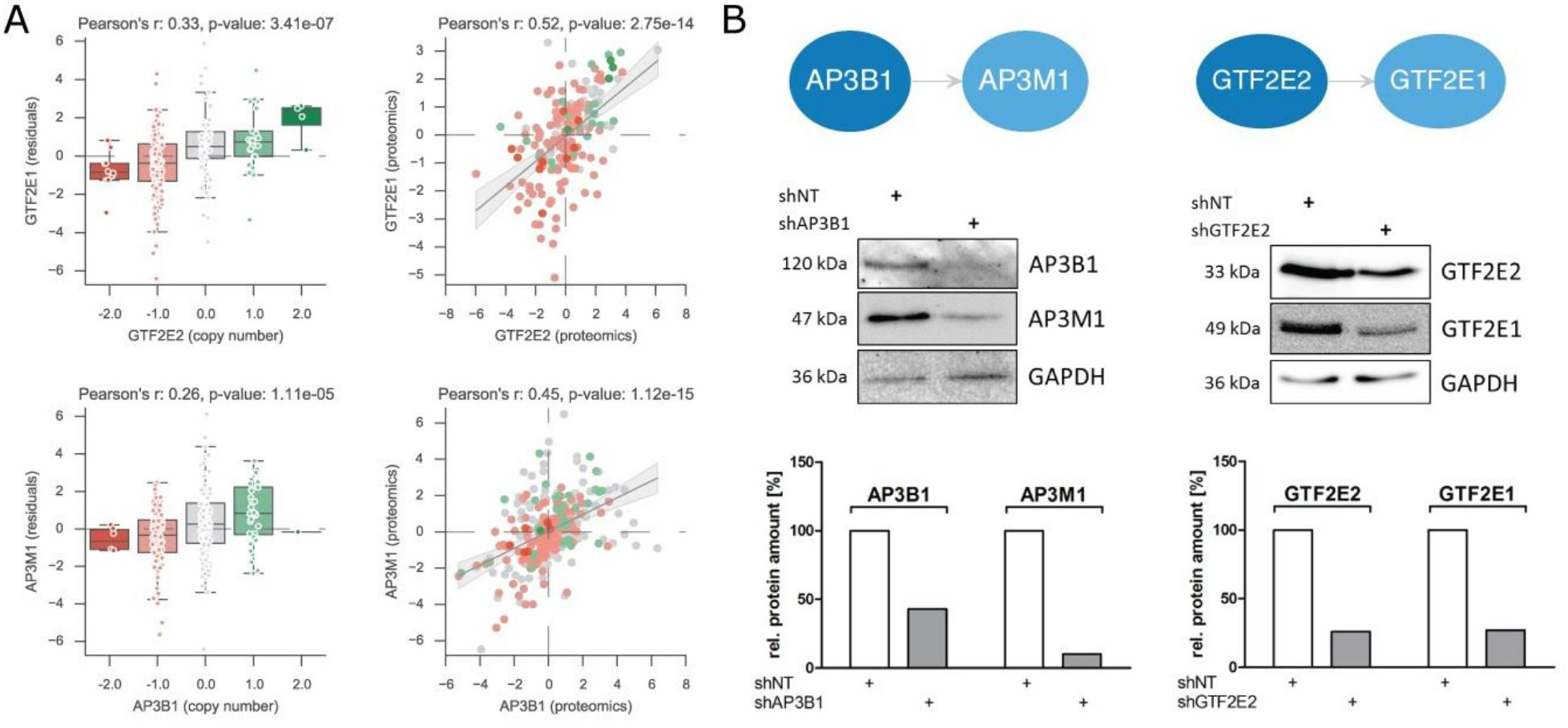
Experimental validation of regulatory interactions among protein complex subunits. A) Correlation of the copy-number profile of the regulatory protein with the protein residuals of the regulated protein (left plot) and agreement at the protein level between the two proteins (right plot). B) Regulatory interactions within the adaptor protein complex 3 (AP3) and the Transcription initiation factor IIE (TFIIE) complexes. shRNA knockdown of the regulatory proteins, AP3B1 and GTF2E2, show strong decrease in the protein abundance of the regulated proteins, AP3M1 and GTF2E1, respectively. Protein abundance changes are measured and quantified by western blot using antibodies specific for the corresponding proteins. The quantified bands in the shAP3B1 and shGTF2E2 experiments were scored relative to the control shRNA (shNT). GAPDH was used as a loading control.

Besides providing us with clues on the assembly pathways of protein complexes, these associations provide concrete examples of trans protein quantitative loci associations (pQTLs) (Chick et al., 2016), illustrating how the genomic variation can have consequences through protein interaction partners.

### Protein complex amplifications associated with high attenuation potential

Having assessed the attenuation of CNVs effects in the proteome we set out to quantify the extent of this regulation in each tumour sample. We reasoned that by stratifying the samples by their capacity to attenuate the CNV changes, we could identify the underlying attenuation mechanisms. Similarly to the protein analysis (Figure 1D) we performed a correlation analysis between the CNVs and transcriptomics and proteomics for each sample (Figure 6A), instead of each protein. Furthermore, recurring to a gaussian mixture model we classified 50 samples (18%) as those having a general strong attenuation effect (see Methods). Such tumour samples have a higher number of genes with strong attenuation, suggesting either an overall increase in degradation or decrease in translation rates in these samples. We then searched for complexes and complex subunits that are more likely to be amplified in the tumours with stronger attenuation and could therefore contribute to the attenuation potential (see Methods). Tumours with strong attenuation effects displayed a significant enrichment of gene amplifications in several complex subunits including genes involved in the endoplasmic reticulum-associated degradation (ERAD) pathway (DERL1 and VIMP), cell polarity (SCRIB, LLGL2 and VANGL2), GPI-anchor biosynthesis (PIGT and PIGU) and RNA interference (AGO2) (Figure 6B and 6C). SCRIB protein complexes have been previously reported to play an important role in cancer progression in breast cancer and their inhibition has been linked to a decrease in cell migration (Anastas et al., 2012). The proteasome system is important for the regulation of focal adhesions in migrating cells (Teckchandani and Cooper, 2016) and inhibition of the proteasome inhibits migration and invasion in breast cancer cells (Xie et al., 2009). However, it is not clear how the overexpression of these cell polarity factors would result in an increase in attenuation potential. Interestingly and consistently with the increased protein attenuation profile of these tumours, we observe amplifications of the ERAD components DERL1 and VIMP, that are part of an endoplasmatic reticulum (ER) complex which is responsible for the retrotranslocation of misfolded proteins to the cytosol for proteasomal degradation (Lilley and Ploegh, 2004; Ye et al., 2004). The association between increased attenuation and amplification of AGO2 could be explained by its role in repressing the initiation of mRNA translation (Kiriakidou et al., 2007).

**Figure 6.**
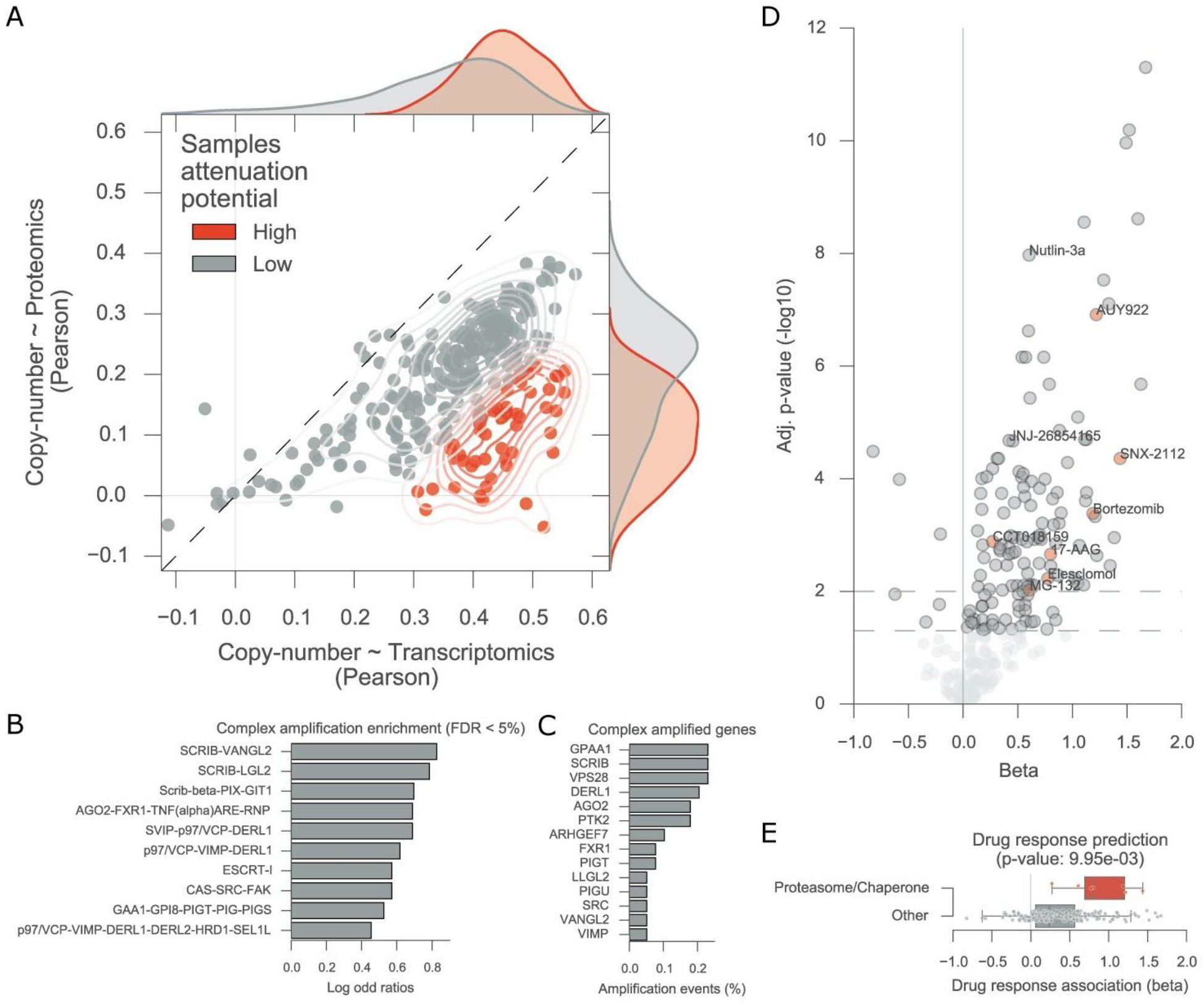
Putative mechanisms for tumour attenuation potential and its association with chaperone/proteasome drug resistance. A) Tumour sample correlations of the copy-number changes and the transcript (x axis) and protein (y axis) measurements. Samples classified with high attenuation potential, in red, display stronger attenuation of the copy-number variation. B) Protein complexes significantly enriched for gene amplifications (FDR < 5 %) on the samples with high protein attenuation. C) Top strongly amplified genes within the significantly enriched complexes. D) Drug response associations performed in a large cell line panel using the cell lines putative attenuation potential as the predictive feature. Significant associations (FDR < 5 %) of chaperone and proteasome inhibitors are labelled and marked in red. Ubiquitin-protein ligase MDM2 inhibitors are labelled. E) Boxplots representing the distributions of the drug associations effect sizes of all the proteasome and chaperones inhibitors in the drug panel.

### Gene expression profile of protein attenuation is associated with specific drug responses

Since the tumours with strong attenuation of CNVs effects displayed particular characteristics, we defined a gene expression signature by systematically correlating each gene with the attenuation potential (see Methods). This signature provides a putative ranking of the agreement between gene expression and the protein attenuation profile of the samples. Next, we explored the capacity of this signature to identify particular cellular states that can be informative for drug response. Samples with a strong correlation with the signature would be predicted to have higher attenuation and could, for example, display a higher proteasomal capacity. Thus, we considered an independent cell line panel for which gene expression and drug response is available (Iorio et al., 2016b), and ranked the cell lines according to their predicted protein attenuation potential (see Methods). Then we assessed the association between this predicted attenuation potential and drug response measurements for 265 compounds (see Methods) (Figure 6D and 6E). Among the top predicted compounds are a proteasome (Bortezomib and MG-132) and chaperone inhibitors (AUY922, 17-AAG, Elesclomol, CCT018159 and SNX-2112) which displayed a significant (FDR < 5%) positive association, suggesting that a stronger predicted attenuation potential is associated with increased resistance to proteasome/chaperone inhibitors (Supplementary Table 4). This unbiased search also revealed significantly positive associations of Nutlin-3a and JNJ-26854165 and the proteome attenuation profile (pearson’s r 0.20 and 0.16, respectively). Interestingly, both compounds target the oncoprotein E3 ligase murine double minute 2 (MDM2) which, in p53 wild type tumours, suppresses the activity of p53 by ubiquitination and thereby is a potential therapeutical target (Shangary and Wang, 2008). The protein attenuation potential predicted for the cell lines also displayed tissue specificity, supporting the idea that proteasomal capacity is constrained by the tissue of origin. This analysis suggests that the gene expression signature for the proteome attenuation may be associated with an increased capacity of the protein quality control machinery and an increased resistance to drugs that target this system.

## Discussion

Although chromosomal rearrangements is an ubiquitous feature of cancer cells, still little is known about their impact on the cellular phenotype, in particular on the regulation of the proteome. Recent proteogenomics studies released by the CPTAC consortium (Cancer Genome Atlas Research Network et al., 2013; Mertins et al., 2016; Zhang et al., 2014, 2016) presented for the first time a broad compendium of proteogenomics measurements across tumours, enabling a systematic analysis of this question.

In this study, we observed that while CNVs have a good agreement with transcript measurements, 23-33% of the proteins undergo post-transcriptional regulation which attenuates the impact of CNVs (Figure 1 C, D). This information alone is very relevant for the identification of causal genes within amplified genome regions. Since amplifications of the attenuated genes are not observed at the protein level, these are less likely to be drivers of cancer progression and similarly less likely to explain changes in drug associations. Notably, this attenuation was more pronounced in protein subunits and complexes, in agreement with previous observations (Dephoure et al., 2014; Stingele et al., 2012). This is likely explained by the fact that the stoichiometry of complexes need to be preserved and that proteins over-represented compared to other members of the complex are likely degraded due to increased instability (McShane et al., 2016). Furthermore, we observed that proteins with stronger attenuation are more quickly ubiquitinated (Kim et al., 2011) (Figure 2D) suggesting that the attenuation may be mostly driven by changes in degradation instead of translation rates.

Protein complex subunit abundance co-regulation was a general feature and this agreement was not so strikingly visible at the transcript abundance, recapitulating recent findings with the same data-set (Wang et al., 2016). Within these subunits we identified 48 putative-regulators that may act as rate-limiting factors for complex assembly, capable of regulating the abundance and assembly of other complex subunits (Figure 4A). This systematic analysis recapitulated known, COG3 - COG2 and COG3 - COG4, and putative new, COG6 - COG2, regulatory interactions within the conserved oligomeric Golgi (COG) complex, which is located in the golgi apparatus and involved in protein sorting and glycosylation (Smith & Lupashin, 2008). COG3 associations were validated in an independent study where knocking down COG3 leads to the depletion of the interacting proteins (Zolov and Lupashin, 2005). Other putative-regulatory interactions were proposed within complexes with important roles in cancer, such as EIF3A regulation of several members of the eIF3 complex, which plays an important role in translation initiation and regulation of protein synthesis (des Georges et al., 2015; Zhang et al., 2007). Also, a regulatory interaction between RPA2 - RPA3 was reported, these proteins are part of the replication protein A (RPA) complex which is involved in the response to DNA damage and thereby is involved in the control of DNA repair mechanisms and the activation of DNA damage checkpoints (Zou et al., 2006). We have experimentally validated two new regulatory interactions, AP3B1 - AP3M1 and GTF2E2 - GTF2E1, within the AP3 and TFIIE complexes, respectively (Figure 5). We also designed experimental validations for RPA2 - RPA3 and for EIF3A - EIF3E, but knocking down RPA2 or EIF3A proved to be lethal for the transfected HCT116 colon cancer cell lines. Interestingly, potential mutual exclusivity associations were present in much lower numbers. The most compelling negative association was SMARCA2 - SMARCA4, which was supported by current literature where the two are reported to be mutual exclusive ATPases (Karnezis et al., 2016) and paralogs (Ori et al., 2016) within the SWI/SNF complex. Dual deficiency of these proteins induces differentiation from a normal cell to high-grade tumour (Karnezis et al., 2016). These results provide examples and putative mechanistic explanations for how variation in copy number of gene expression of a protein can have trans effects in the abundance of interacting proteins, as seen in protein quantitative trait loci analyses (Chick et al., 2016).

Tumour samples with strong attenuation of CNVs effects in protein abundance displayed a significant enrichment for amplifications of several protein complexes involved in the response to misfolded proteins in the ER, cell polarity, trafficking and gene repression. The amplification of genes associated with cell polarity in cells with increased attenuation potential would suggest that increased cell migration might result in an increased proteasomal function or decreased translation rates. We derived a gene expression signature to characterise the attenuation effect and used this to associate with drug responses in an independent large panel of approximately 1,000 cell lines and 265 compounds (Iorio et al., 2016b) (Figure 6D). Proteome attenuation was associated with increased resistance to proteasome and chaperone inhibitors, suggesting that tumours where attenuation is more pronounced are more resistant to perturbations in the chaperone/proteasome system. Interestingly, the two compounds in the screen targeting MDM2 were among the top associated with the gene expression signature, suggesting that tumours with high predicted attenuation potential may have a high proteasome capacity and therefore be less sensitive to the inhibition of MDM2, that is the E3 ligase responsible for the degradation of p53 in TP53 wild-type tumours (Shangary and Wang, 2008). Additional work will be required to conclusively validate the putative associations between the attenuation potential and the drug responses.

In this study, we provide novel insights into how cancer cells regulate their proteome in the presence of abnormal chromosomal rearrangements, in particular of copy-number alterations. This study emphasises the importance of integrating multiple types of biological data to allow functional assessment of genomic alterations. While relevant insights were possible here, more tailored experimental work should be carried-out to understand the mechanisms of how are the co-regulatory effects of complexes maintained. Mutations can have similar implications to copy-number alterations by affecting the correct synthesis of the protein or its transcript, but these are generally harder to interpret. We have uncovered a potential way in how cancer cells manage to cope with often dramatic chromosomal rearrangements (Thompson and Compton, 2011) and these can possibly provide insights into their functional implications and hopefully open novel therapeutic opportunities.

## Methods

### Data compendium

Proteomics measurements at the protein level for the three tumour types analysed here were compiled from the CPTAC data portal (Edwards et al., 2015) (accession date 2016/07/06) for the following publications: BRCA (Mertins et al., 2016), HGSC (Zhang et al., 2016) and COREAD (Zhang et al., 2014). Transcriptomics RNA-seq raw counts were acquired from (Rahman et al., 2015) (GSE62944) and processed using the Limma R package (Ritchie et al., 2015) with the voom transformation (Law et al., 2014). GISTIC (Mermel et al., 2011) thresholded copy-number variation measurements and clinical data were obtained from the http://firebrowse.org/ portal (accession date 2016/06/08).

### Data processing and normalisation

Transcriptomics raw counts were downloaded from (Rahman et al., 2015) (GSE62944). To ensure that lowly expressed transcripts are removed, genes with average counts per million (CPM) across samples lower or equal to 1 were excluded. Data was normalised by the trimmed mean of M-values (TMM) method (Robinson and Oshlack, 2010) using edgeR (Robinson et al., 2010) R package. Finally, the log-CPM values derived from the voom (Law et al., 2014) function in Limma (Ritchie et al., 2015) package were extracted for this analysis.

Coverage of the proteomics samples was assessed using the jaccard index for each sample with matching transcriptomics. Transcriptomics and proteomics measurements were used at the gene symbol level annotation. For each sample it was only considered transcripts passing the expression threshold, defined above, and proteins with matching measurement. The jaccard index for each sample was calculated with the intersection over the union.

Considering that proteomics and transcriptomics principal component analysis (PCA) revealed associations with possible confounding factors, i.e. age, gender, tumour type and measurement technology, we regressed them out from the original data-sets using linear regression models (Figure S1). For each protein a multiple linear regression model was fitted with protein measurements across the tumour samples as the dependent variable and the confounding factors mentioned above as independent discrete variables, apart from the age which was represented with a continuous variable. Once the model was fitted the estimated weights of the covariates were used to regress-out their impact in the protein measurement and thereby removing their effects (Figure S2). Due to the sparseness of mass-spectrometry measurements for the proteomics data-set we only considered proteins that were consistently measured in at least 50% of the samples, leaving a total of 6,734 proteins. The same procedure was performed in the transcriptomics measurements. Transcript and protein measurements were normalised and centered across the samples using a gaussian kernel density estimation function.

### Proteome attenuation analysis

Agreement between the copy-number variation and the transcriptomics and proteomics was calculated for each gene/protein across the tumour samples using pearson correlation coefficient. Enrichment of biological processes for proteins displaying an attenuation of the correlation at the protein level compared to the transcript level was performed using Gene Set Enrichment Analysis (GSEA) (Subramanian et al., 2005). For the enrichment we used the protein attenuation level, which is calculated by the difference between the pearson coefficient of the transcript correlation (correlation between copy-number variation and transcript measurements) and the pearson coefficient of the protein correlation (correlation between copy-number and protein measurements). To ensure a normal distribution centered around zero for the GSEA enrichments a gaussian kernel density estimation function was used to normalise the protein attenuation distribution. Gene signatures of Gene Ontology (GO) (Ashburner et al., 2000; The Gene Ontology Consortium, 2015) terms for biological processes (BP) and cellular compartments (CC) were acquired from the MSigDB data-base (Subramanian et al., 2005). Gene signatures of post-translational modifications (PTMs) were also used and acquired from Uniprot data-base (The UniProt Consortium, 2015). The estimated enrichment scores were statistically assessed by performing 1,000 random permutations of the signatures and p-values were then adjusted using false-discovery rate (FDR).

Proteins were classified according to their copy-number attenuation effect using a gaussian mixture model with 2 mixture components. Proteins in the group with larger mean attenuation were considered highly attenuated. More stringent classification of the attenuation effect was performed by only considering attenuated proteins with an absolute attenuation score higher than 0.3.

For samples the attenuation potential was estimated similarly as for proteins but instead correlations were calculated across the proteins measured in the sample. Samples were then classified as before with a gaussian mixture model with 2 mixture components. Enrichment analysis of amplifications in protein complexes in tumour samples with high protein attenuation potential was performed using SLAPenrich (Iorio et al., 2016a).

A gene expression signature of the sample attenuation potential was calculated by systematically correlating the samples attenuation potential with each gene in the transcriptomics data-set.

### Pairwise correlation analysis

Correlations between protein pairs, or genes, across samples were calculated using pearson correlation coefficient. Only proteins that were also measured at the transcript level were considered, i.e. 6,434. The systematic analysis of all unique pairwise correlations generated a total of 41,389,922 correlation coefficients both at the protein and gene level.

Protein sets of known protein complexes were acquired from the CORUM data-base (Ruepp et al., 2010, 2008). A protein-protein interaction list of the complexes was assembled by considering that two proteins interact if they are present within the same complex at least once, this generated a total of 67,927 interactions. Indirect but functional associations were also considered by using the STRING data-base (Franceschini et al., 2013). For STRING only interactions with the highest confidence score (900) were used performing a total of 214,815 interactions. 9,273 protein interactions within signalling pathways were assembled from kinase/phosphatase-substrate interactions reported in SIGNOR data-base (Perfetto et al., 2016). Metabolic enzyme interactions associated with metabolic pathways were extracted from KEGG pathways (Kanehisa et al., 2016) reported in MSigDB (Subramanian et al., 2005). Two enzymes were considered to be interacting if they were present in the same metabolic pathway, making a total of 121,134 interactions. Enrichment of the different types of protein-protein interactions, i.e. complexes, functional, signalling and metabolic, were estimated using receiving operating characteristic (ROC) curves and by calculating the area under the ROC curve (AROC). True-positive sets of protein interactions were defined as the ones reported in the different resources used. Due to the strong unbalance between the number of true positives and false positives the ROC curves were calculated using 5 different and randomised sets within the false positive group. The variability of the AROC score is represented by error bars in Figure 3C.

### Proteogenomics analysis to identify protein complex regulators

The identification of protein complex regulators only focused on protein-protein interactions reported in the CORUM data-base (Ruepp et al., 2010, 2008) with a protein-protein interaction list assembled as described before.

For each protein-protein interaction reported within a complex, its association was tested using two linear regression models. Given a pair Protein Y ~ Protein X (Py ~ Px), a first linear model is used to regress-out the transcript variability from the protein measurement of Py. The dependent variable of the model is the proteomics measurements of Py and the independent variable is the transcriptomics measurements (Ty) (Eq. 1):

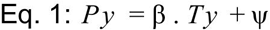

The model is fit with an intercept (for simplicity omitted from Eq.1) and noise term, ψ. After fitting the estimated weight (β), the residuals of Py (Py’) are calculated as (Eq. 2):

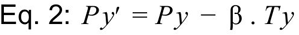

Py’ represents the variability measured due to post-transcriptional and post-translational regulation. Then a second linear model is performed to calculate the association between Py’ and the CNV of Px, (Px) (Eq: 3):

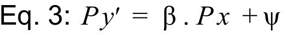

Statistical significance is estimated by calculating an F statistic over an F-distribution, p-values are then adjusted using FDR correction. A total of 58,627 tests are performed. The same analysis is performed using transcriptomics measurements instead of the copy-number of Px, generating a total of 57,462 tests. Associations estimated with the copy-number variation that are significant with the transcriptomics are highlighted with a red border in Figure 3A.

### Cell lines drug response analysis

Gene expression measurements (E-MTAB-3610) acquired with Affymetrix Human Genome U219 array for approximately 1,000 cell lines was used in this analysis (Iorio et al., 2016b). Drug response measurements were obtained as the area under the curve (AUC) for 265 compounds (Iorio et al., 2016b). Cell lines proteome attenuation potential was calculated by performing pearson correlation between their transcriptomics profile and the proteome attenuation potential signature derived from tumours. Cell line correlations with the signature were then used as a feature in single linear regression models to systematically predict the response of each compound in the screen.

### Cell culture and sample collection

The human colon cancer cell line HCT116 was cultivated in McCoys 5a medium supplemented with 10% FBS and 1% penicillin/streptomycin under standard culture condition. AP3B1 and GTF2E2 silencing was obtained by lentiviral short hairpin RNA (shRNA) delivery. shNT (“non target”) clones were used as control cells. For protein sample isolation, 10^6^ cells of shNT, shAP3B1 or shGTF2E2 clones were plated in 10 cm culture dishes for 48h. Afterwards cells were lysed in RIPA buffer to obtain total protein samples. Protein content was determined by DC™ protein assay as recommended by the manufacturer (Bio-Rad laboratories Inc, Cat.#: 500-0116, Hercules, CA USA).

### shRNA delivery via lentiviral transduction

The applied shRNA plasmids are part of the MISSION^®^ shRNA product line of Sigma Aldrich (shAP3B1, Cat.#: SHCLND-NM_003664, TRC clone: TRCN0000286136; shGTF2E2, Cat.#: SHCLND-NM_002095, TRC clone: TRCN0000020775; shNT, Cat.#: SHC016-1EA) and were delivered via lentiviral transduction using a second generation lentiviral packaging system. Therefore, HEK293T cells were co-transfected with the appropriate pLKO.1 transfer-vector (shRNA containing vector), psPAX2 (the packaging vector) and pMD2. G (the vector that encodes for the viral envelope protein) using jetPEI transfection reagent according to manufacturer’s recommendation (Polyplus transfection, Cat.#: 101-10N, Illkirch, France). Virus-containing supernatants were used for cell transduction.

### Western Blot validation

Predicted protein complex formations of AP3B1_AP3M1 and GTF2E2_GTF2E1 were validated by western blot technique. Total protein lysates (30 µg) were heat-denatured in NuPAGE LDS sample buffer containing dithiothreitol (Thermo Scientific, Cat.#: NP0008, Waltham, MA USA) and loaded on 12% denaturing polyacrylamide gels for separation. SDS-PAGE was conducted with a 2-Step protocol (Step1: 20min 50V constant, Step2: 120min 120V constant). Proteins were transferred to nitrocellulose membranes by tank-blotting (140min at 70V constant). Afterwards membranes were blocked with 5% milk (MP) in TBS-T. All washing steps were conducted with TBS-T. Membranes were incubated with primary antibodies mc mouse α-AP3B1 (abnova, Cat.#: H00008546-B01P, Taipei City, Taiwan; 1:500), mc rabbit α-AP3M1 (abcam, Cat.#: ab201227, Cambridge, UK; 1:1000), mc rabbit α-GTF2E2/TFIIEbeta (abcam, Cat.#: ab187143, Cambridge, UK; 1:10000) or mc rabbit α-GTF2E1/TFIIEalpha (abcam, Cat.#: ab140634, Cambridge, UK; 1:1000) overnight at 4°C. Protein expression of GAPDH was used as loading control using α-GAPDH(D15H11) antibody (CST, Cat.#: 5174S, Cambridge, UK; 1:2000). All primary antibodies were diluted in 5% MP TBS-T. Secondary antibodies used in this work are: HRP-conjugated anti-rabbit IgG (CST, Cat.#: 7074S, Cambridge, UK) for the detection of AP3M1 (1:2000), GTF2E2 (1:1000), GTF2E1 (1:2000) & GAPDH (1:2000), and HRP-conjugated anti-mouse IgG (CST, Cat.#: 7076S, Cambridge, UK) for the detection of AP3B1 (1:5000). Secondary antibodies were diluted in TBS-T and incubated for 1h at room temperature. Quantity One^®^ software (Bio-Rad laboratories Inc., Hercules, CA USA) was used for densitometry.

**Supplementary Table 1.** Overview of the biological measurements available for the TCGA and CPTAC samples.

**Supplementary Table 2.** Pearson correlation coefficients for all genes overlapping between copy-number variation, transcriptomics and proteomics data-sets.

**Supplementary Table 3.** Protein complex regulatory associations estimated between the putative regulatory protein, Px, and the regulated protein, Py, using the copy-number variation and transcriptomics measurements of Px.

**Supplementary Table 4.** Tumour protein attenuation potential gene-expression signature and drug response analysis results.

### Code availability

All the computational analyses were performed in Python version 2.7.10, apart from the transcriptomics RNA-seq processing which as done in R version 3.3.1 with Limma package version 3.28.21 and edgeR 3.14.0, and are available under GNU General Public License V3 in a GitHub project in the following url https://github.com/saezlab/protein_attenuation. Plotting was done using Python modules Matplotlib version 1.4.3 (Hunter, 2007) and Seaborn version 0.7.0 (Waskom et al., 2014). Generalised linear models were built using Python module Sklearn version 0.17.1 (Pedregosa et al., 2011). Data analysis and structuring was carried out using Python module Pandas version 0.18.1 (McKinney and Others, 2010).

## Author Contributions

J. S. R and P. B. conceived and led the study. E. G. carried out the analysis. A. F. and T. C. designed the experimental validations. A. F. carried out cell cultures and knocking-down experiments. L. G. A. contributed to the analysis. E. G., J. S. R. and P. B. wrote the paper.

## Acknowledgments

We thank Michael Schubert for help integrating the copy-number variation data. We gratefully acknowledge helpful comments from Colm Ryan, Marc Brehme, David Ochoa, Romain Studer, Haruna Imamura and Theodoros Roumeliotis.

---

**Supplementary Figure 1.**
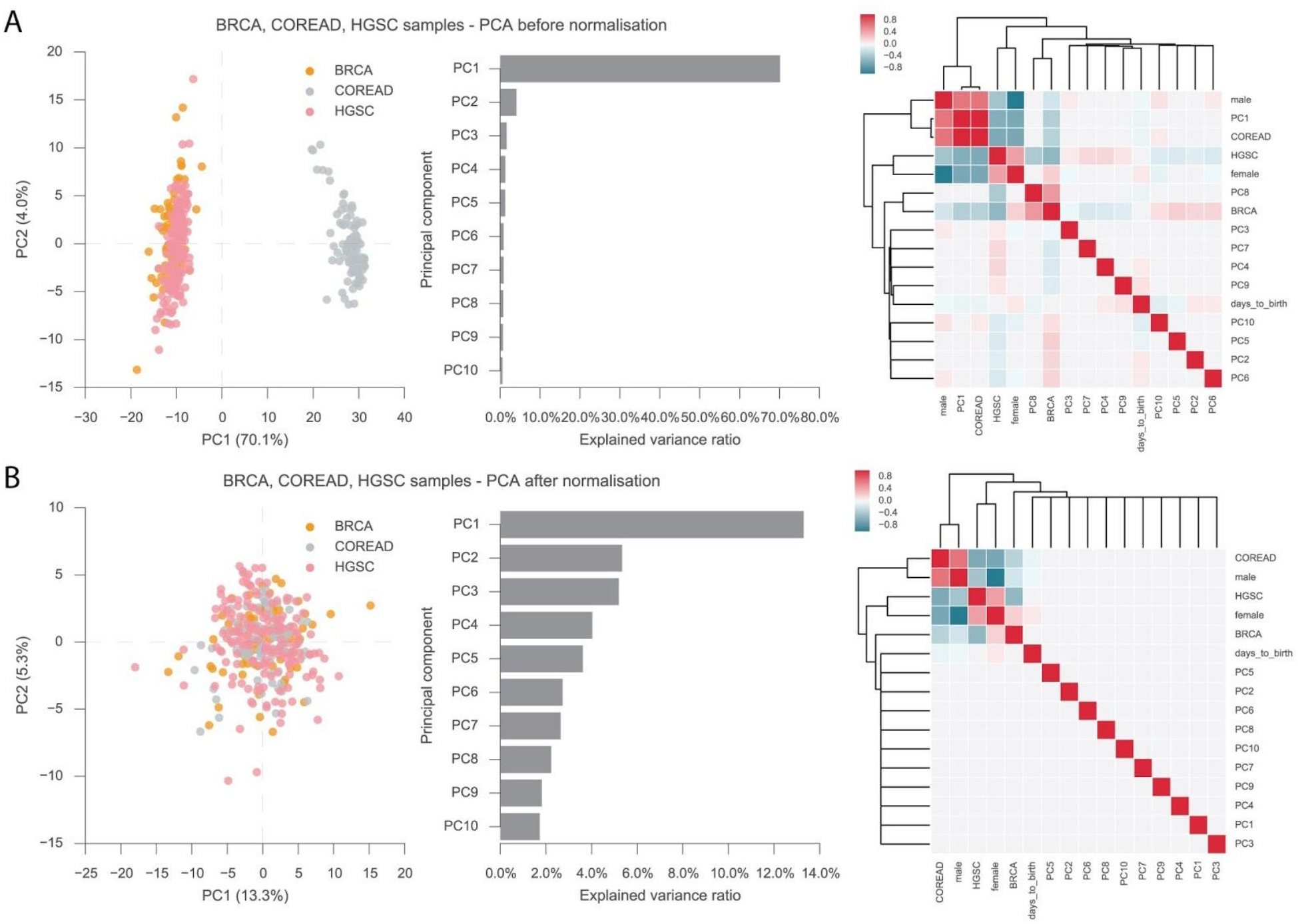
Proteomics data-sets PCA analysis using proteins consistently measured across all the samples and pearson correlation coefficient between the first 10 principal components and the possible confounding factors, i.e. age, tumour type and gender. A) Analysis performed on the original proteomics data-sets. B) Analysis performed on the proteomics data-set after the confounding factors were regressed-out.

**Supplementary Figure 2.**
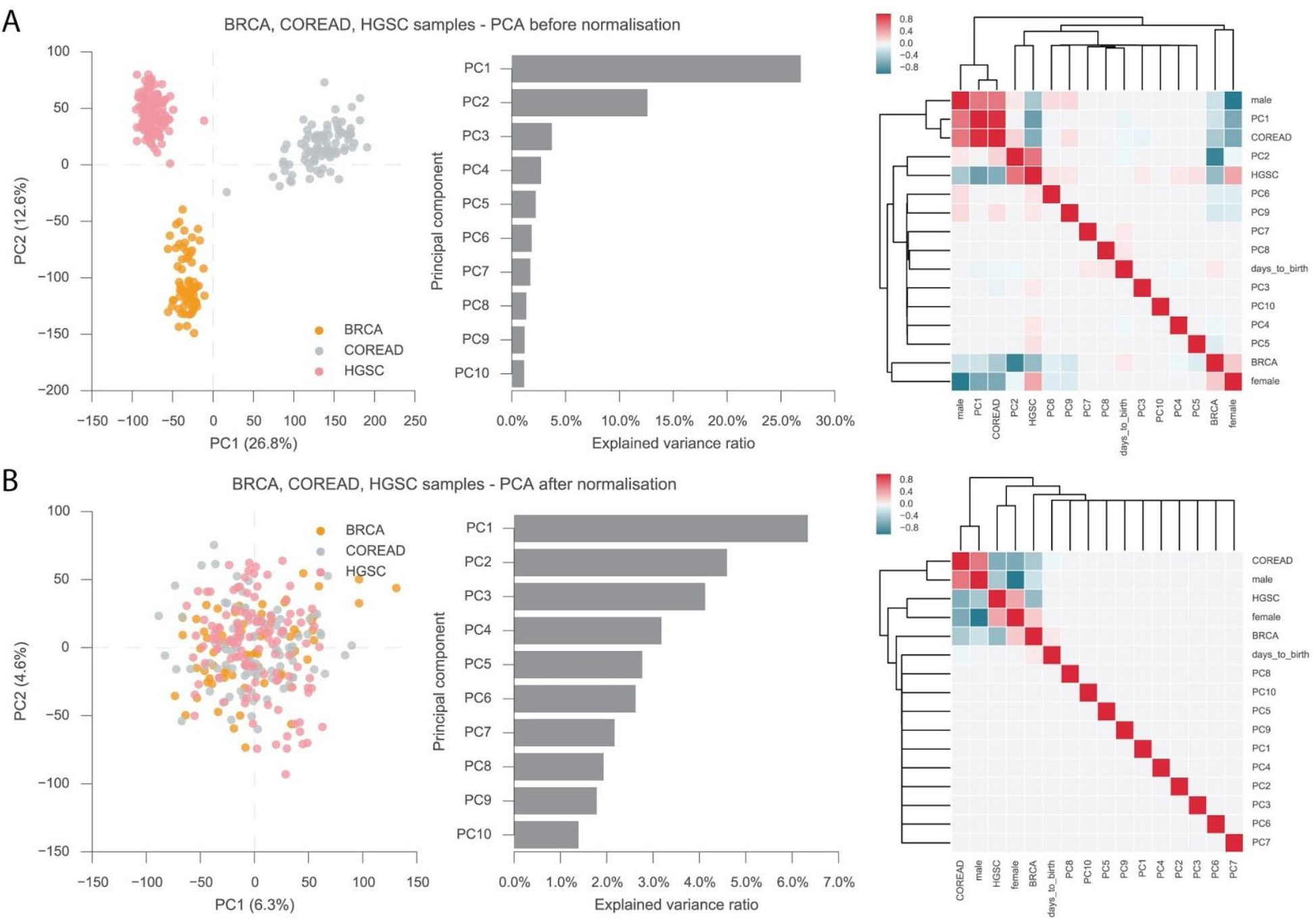
Transcriptomics data-sets PCA analysis and pearson correlation coefficient between the first 10 principal components and the possible confounding factors, i.e. age, tumour type and gender. A) Analysis performed on the voom transformed transcriptomics measurements. B) Analysis performed on the transcriptomics data-set after the confounding factors were regressed-out.

**Supplementary Figure 3.**
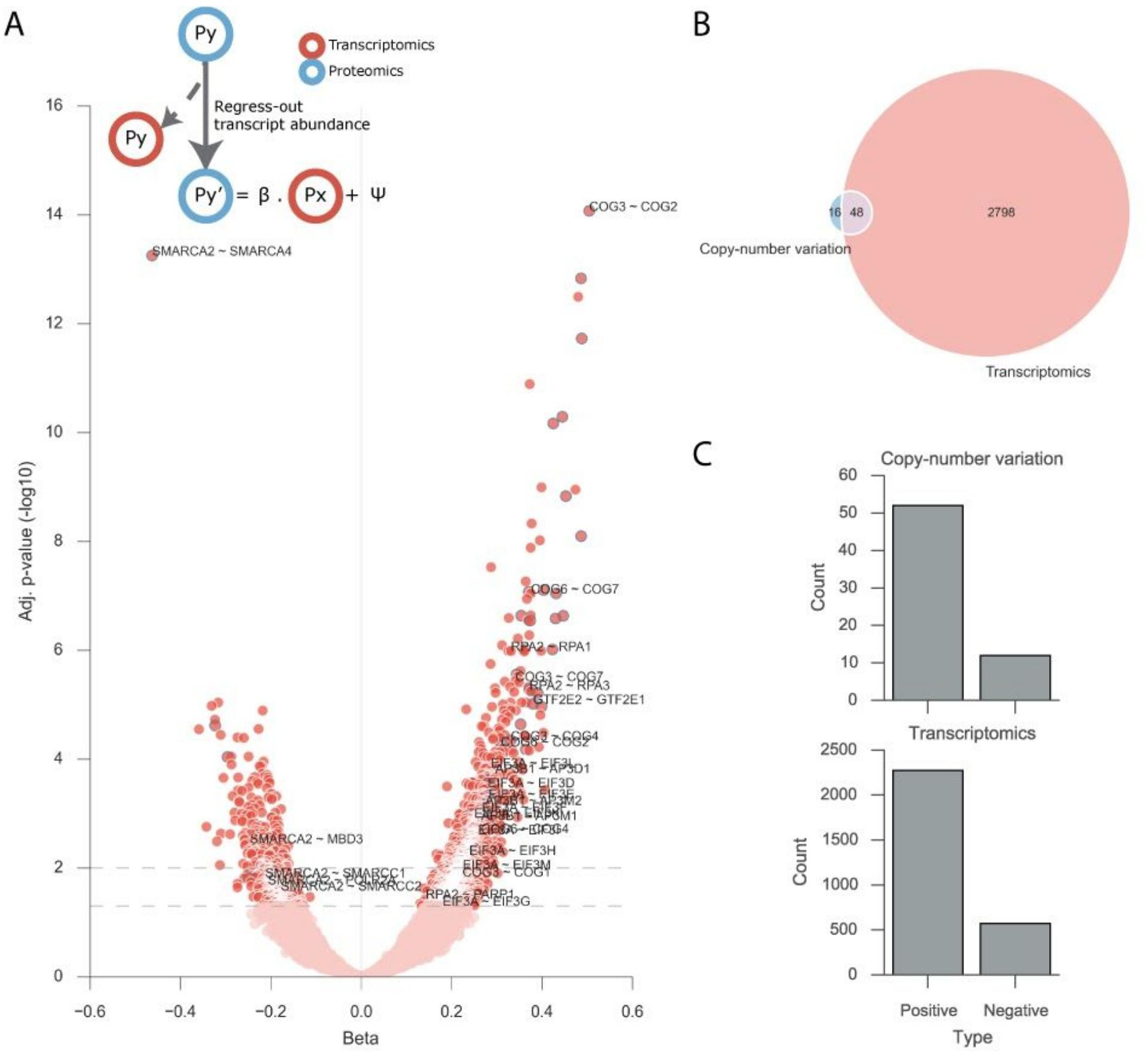
Protein complexes regulatory interactions identified using transcriptomics of the putative regulatory protein (Px). A) Volcano plot representing the effect size on the x axis and FDR adjusted p-value on the y axis. Diagram representing the linear model used to perform the associations. B) Overlap between the significant regulatory associations found using the copy-number variation and transcriptomics of the Px proteins. C) Number of significant associations with a Positive or Negative effect size.

